# Investigating the antioxidant effects of *Aloe vera*: Possible role in regulating lipid profile and liver function in high fat and fructose diet (HFFD) fed mice

**DOI:** 10.1101/2022.04.20.488905

**Authors:** Abdullahi Mukhtar Abubakar, Nathan Isaac Dibal, Martha Orendu Oche Attah, Samaila Musa Chiroma

## Abstract

Fat rich diets are believed to induce obesity and contributes to the development of diabetes and cardiovascular disease while high fructose diet was reported to increase gut surface area and enhance nutrient uptake resulting in weight gain. The study investigate the role of *Aloe vera* supplementation on lipid profiles, oxidative stress as well as liver and hear histology in high fat and fructose diet fed mice. Twenty mice were distributed into four groups (n=5). The groups received regular diet, high fat and fructose died (HFFD), HFFD plus 10% *Aloe vera* (HFFD+AV1) and HFFD plus 20% *Aloe vera* (HFFD+AV2) respectively for 10 weeks. The cholesterol level of HFFD+AV treated mice were significantly lower compared to HFFD treated mice. The ALT level was significantly increased in HFFD treated mice relative to the control. *Aloe vera* significantly improve albumin level as well as Catalase and superoxide dismutase activities of HFFD treated mice. The liver tissues of control and HFFD+AV2 treated mice showed normal hepatocytes. The study suggest that *Aloe vera* supplementation could protect against HFFD induced oxidative stress and hyperlipidemia. These findings might be used for further research on food supplementation for the control of metabolic disorders.

## Introduction

Obesity and hyperglycemia have been of public health concern due to the associated complications, increasing incidence, high cost of management/treatment and high mortality rate [1]. Obesity was reported to be associated with liver disease, hyperglycemia, oxidative stress and hypercholesterolemia [2]. A strong correlation between obesity and hyperglycemia has been reported in previous studies and it often results in fatty liver disease [3]. Non-alcoholic fatty liver disease (NAFLD) is characterized by steatosis and about five per increase in triglycerides with no evidence of alcohol abuse [4]. Obesity and diabetes are the most common risk factors of NAFLD. NAFLD was reported to occur in more than 50% of overweight and diabetic individuals [5]. NAFLD is believed to progress to non-alcoholic steato-hepatitis (NASH) in about 20% of patients; the rate of progression from NAFLD to NASH is high in obese, hypertensive and dyslipidemia individuals [6]. While fat rich diets are believed to induce obesity and contributes to the development of diabetes and cardiovascular disease [7], high fructose diet was reported to increase gut surface area and enhance nutrient (fat) uptake resulting in weight gain [8].

*Aloe vera* is a drought resistant succulent tropical xerophyte belonging to the Liliaceous family with many pharmacological properties [9]. Earlier report suggest that *Aloe vera* have anti-hyperglycemic and antioxidant properties with the potentials of ameliorating lipid accumulation and obesity [10]. The plant *Aloe vera* have been used in ancient Egyptian and Chinese medicine to treat fever, burns and wounds [11]. It contains many bioactive compounds including enzymes, vitamins, anthroquinones and amino acids with diverse medicinal properties ranging from antiseptic, anti-obesity, anti-inflammatory, antioxidant to antibacterial effects [12–14]. With these numerous health benefits of *Aloe vera,* the current study aimed to investigate the role of *Aloe vera* supplementation on lipid profiles, oxidative stress as well as liver and hear histology in high fat and fructose diet fed mice.

## Materials and Methods

### Diet formulation

*Aloe vera* were collected from a garden in Maiduguri, Nigeria and identified by a botanist in the Faculty of Pharmacy, University of Maiduguri herbarium (UMM/FPH/ASH/001). The gel was separated and used for die formulation. Normal rat chow consist of 4% fat, 15% protein and 6% fibre. High fat and fructose diet (HFFD) consist of 70% normal chow with 30% margarine and 15% fructose in drinking water. *Aloe vera* supplementation was formulated in two different regiment; 90g of HFFD plus 10g *Aloe vera* (HFFD+AV1) and 80g of HFFD plus 20g Aloe vera (HFFD+AV2) respectively.

### Animal treatment and Ethics approval

Twenty (20) male six weeks old BALB/c mice (18-21g) were purchased from the National Veterinary Research Institute (NVRI) Vom, Nigeria and kept in the Animal house Department of Biochemistry, University of Maiduguri to acclimatize. The research was approved by Department of Human Anatomy Ethical committee, University of Maiduguri (UM/HA/PGR20.21-08800). The study was carried out in accordance with the ARRIVE guidelines and the National Institute of Health Guide for the Care and Use of laboratory Animals. All surgical procedures were performed under ketamine hypochlorite Anesthesia and efforts were made to minimize suffering.

### Experimental design

Twenty mice were randomly distributed into four groups (n=5). The groups received normal chow, HFFD, HFFD+AV1 and HFFD+AV2 respectively for 10 weeks. The mice in each group were marked for easy identification. All the mice were euthanized thereafter, blood samples were collected and centrifuged at 5000 rpm for 10 minutes. The liver was dissected, fixed in 10% formalin, and processed for light microscopy.

### Biochemical parameters

The serum levels of aspartate aminotransferase (AST), alkaline phosphatase (ALP), albumin, alanine aminotransferase (ALT), total protein concentration, triglycerides, cholesterol, high density lipoprotein (HDL) and low density lipoprotein (LDL) were estimated from 4 four mice in each group using enzyme linked immune-sorbet assay (ELISA) kit (NeoScientific, USA).

### Antioxidant activity

The activities of catalase (CAT), reduced glutathione (GSH), superoxide dismutase (SOD) and Malondialdehyde were evaluated from liver homogenate of four mice in each group as described by Aebi [15], Rajagopalan et al. [16], Fridovich [17] and Akanji et al. [18] respectively.

### Histological study

The fixed liver tissues were dehydrated in graded alcohol, cleared in xylene, embedded in paraffin and sectioned at 5μm. Tissue sections were stained with heamatoxylin and eosin (H&E) and micrographs were taken at x200 magnifications using digital microscope camera (AmScope, UK).

### Statistical analysis

GraphPad prism 7 (GraphPad, USA) was used to analyzed the data. One-way analysis of variance and Tukey post-hoc test was carried out and the results were expressed as Mean ± standard error of mean (SEM). P<0.05 was considered statistically significant.

## Results

### Lipid profile

Cholesterol level was significantly increased (P<0.05) in HFFD treated mice relative to the control. On the other hand, the cholesterol level of HFFD+AV treated mice were significantly lower (P<0.05) compared to HFFD treated mice (Fig 1). High density lipoprotein level was significantly (P<0.05) reduced in HFFD and HFFD+AV treated mice compare to control. However, low density lipoprotein (LDL) level of HFFD treated mice were significantly higher (P<0.05) compared to the control and HFFD+AV treated mice. HFFD+AV treated mice had significantly lower (P<0.05) LDL level compare to HFFD treated mice (Fig 1).

**Fig 1.**
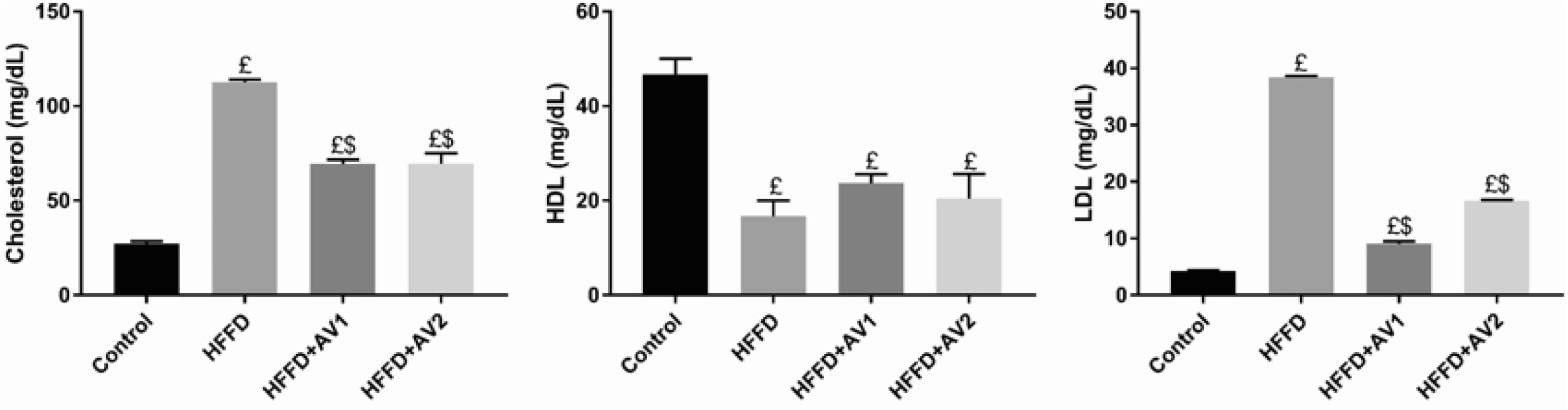
Lipid profile of HFFD and HFFD+AV treated mice. £ and $ indicates significant difference with control and HFFD at (P<0.05). HFFD=High fat and fructose diet. HFFD+AV1= 90g of HFFD plus 10g *Aloe vera.* HFFD+AV2= 80g of HFFD plus 20g *Aloe vera,* n=4

### Liver function

The level of ALT was significantly increased (P<0.05) in HFFD treated mice relative to the control. Nonetheless, ALT level of HFFD+AV treated mice was significantly lower compared to HFFD and HFFD+AV treated mice (Fig 2). The levels of AST and ALP were not significantly changed (P>0.05) in HFFD and HFFD+AV treated mice compared to the control. Albumin and total protein concentration of HFFD treated mice were significantly reduced (P<0.05) relative to the control and HFFD+AV treated mice. Nevertheless, albumin and total protein concentrations were significantly higher (P<0.05) in HFFD+AV2 treated mice relative to the control (Fig 2).

**Fig 2.**
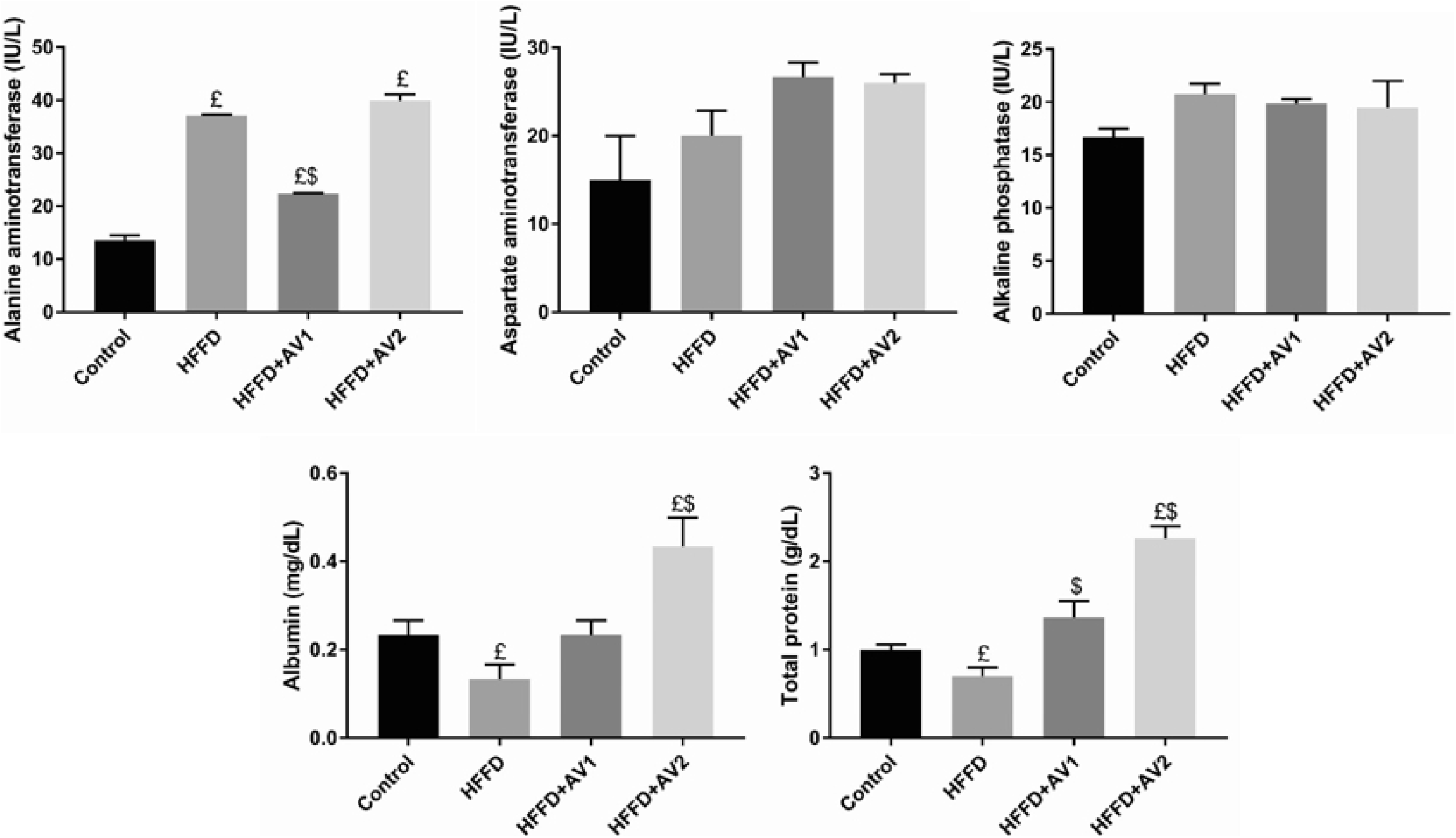
Liver marker enzymes of HFFD and HFFD+AV treated mice. £ and $ indicates significant difference with control and HFFD at (P<0.05). HFFD=High fat and fructose diet. HFFD+AV1= 90g of HFFD plus 10g *Aloe vera.* HFFD+AV2= 80g of HFFD plus 20g *Aloe vera*, n=4

### Antioxidant activity

Catalase and superoxide dismutase activities of HFFD treated mice were significantly lower (P<0.05) compared to the control and HFFD+AV treated mice. However, their activities were not significantly changed (P>0.05) in HFFD+AV treated mice relative to the control (Fig 3). A dose dependent non-significant increase (P>0.05) in GSH activity was observed in HFFD+AV treated mice relative to the control and HFFD treated mice. Also, a non-significant change (P>0.05) in MDA level was observed in HFFD treated mice compared to the control and HFFD+AV treated mice (Fig 3).

**Fig 3.**
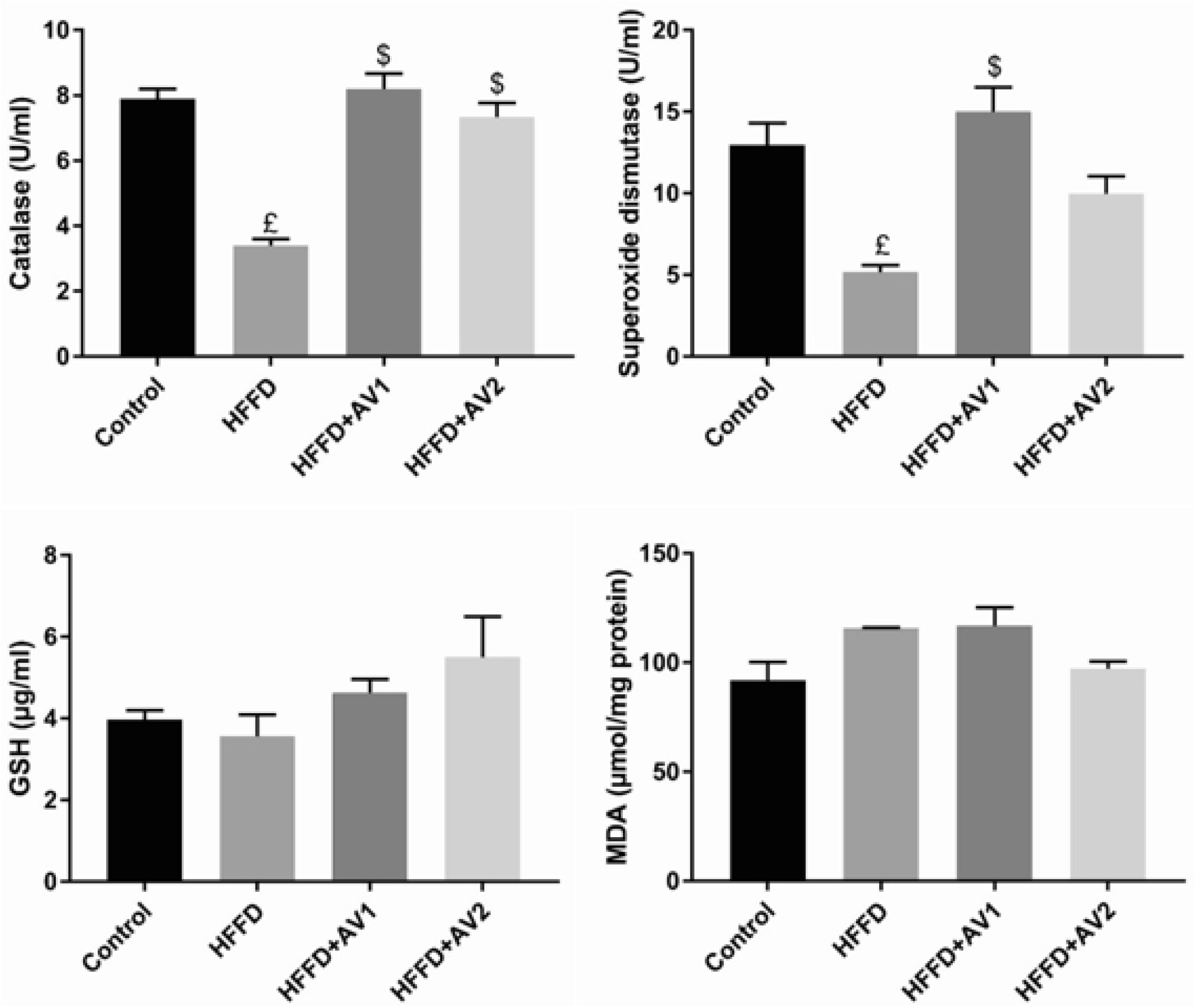
Oxidative stress markers of HFFD and HFFD+AV treated mice. £ and $ indicates significant difference with control and HFFD at (P<0.05). HFFD=High fat and fructose diet. HFFD+AV1= 90g of HFFD plus 10g *Aloe vera.* HFFD+AV2= 80g of HFFD plus 20g *Aloe vera,* n=4

### Histological study

The liver tissues of control and HFFD+AV2 treated mice highlighted normal hepatocytes. However, the liver of HFFD and HFFD+AV1 treated mice showed numerous hepatic vacuoles suggesting fat droplet within the liver tissues (Fig 4).

**Fig 4.**
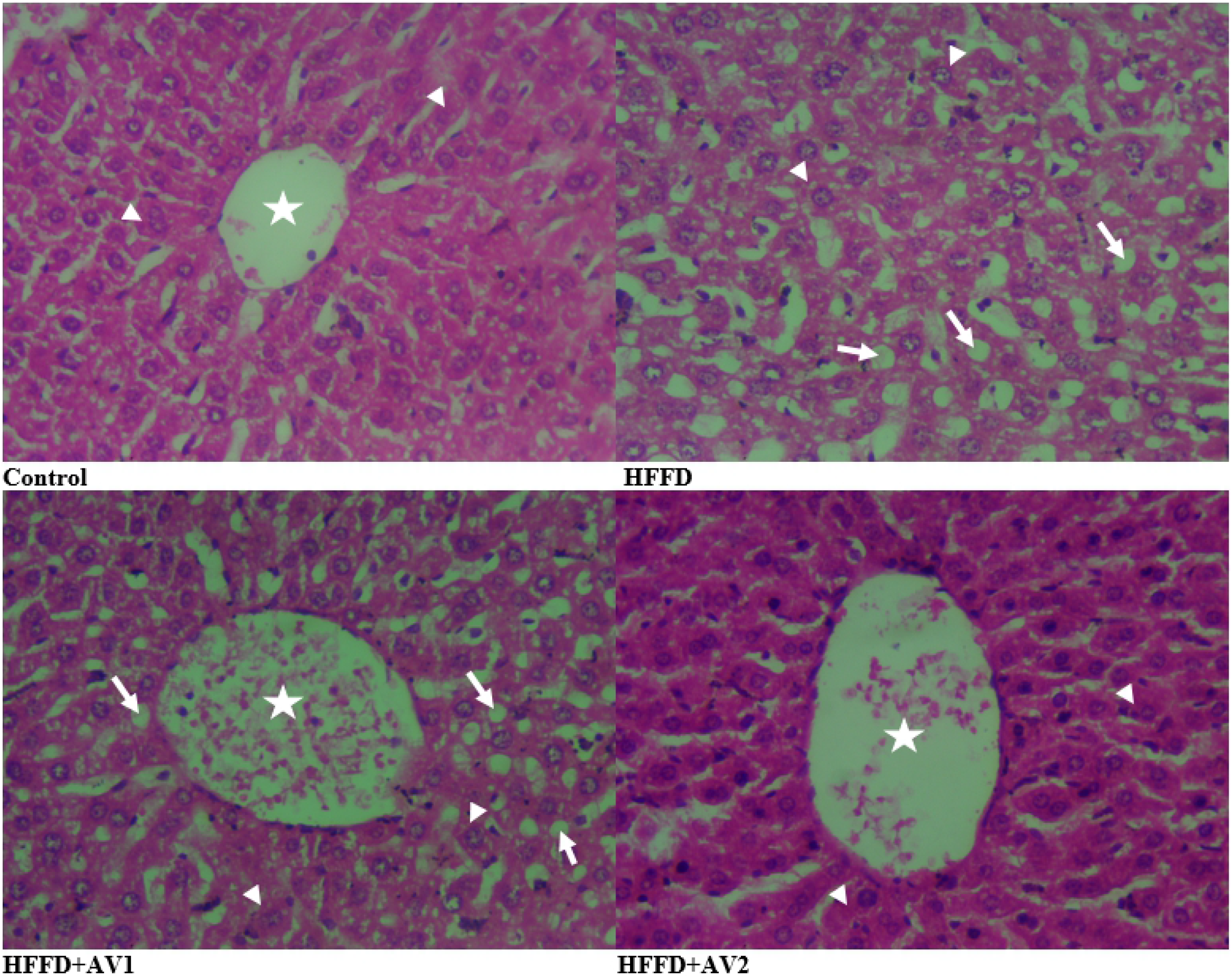
The liver tissue of HFFD and HFFD+AV treated mice. H&E x400 magnification. HFFD=High fat and fructose diet. HFFD+AV1= 90g of HFFD plus 10g *Aloe vera.* HFFD+AV2= 80g of HFFD plus 20g *Aloe vera*.

## Discussion

Several literatures demonstrated a link between fat rich and high fructose diet with dyslipidemia [19–20]. Hence, the high serum levels of cholesterol and LDL with low HDL level that was observed on HFFD fed mice in the present study. Dyslipidemia is induced by fat rich and high fructose diet either through liver synthesis of very low density lipoprotein when free fatty acid and/or triglycerides gets to the liver or through the increase in gut surface area to enhance fat uptake following high fructose diet intake [8,21]. In the present, *Aloe vera* prevents dyslipidemia in HFFD fed mice. This might be that, *Aloe vera* prevents very low density lipoprotein synthesis in the liver or reduce suppress fat uptake from the gut. *Aloe vera* was reported to promote lipolysis thereby preventing dyslipidemia and obesity related complications [22]. The resultant effect of *Aloe vera* might be the cause of normal liver tissue of HFFD+AV2 treated mice in the present study. The mechanism which *Aloe vera* prevents fatty liver might be by increasing lipolysis and hence preventing fat accumulation in the liver.

The present study reported high ALT level and fatty liver in HFFD fed mice. High fat diet is associated with fatty liver and elevated serum levels of liver marker enzymes [23,24]. *Aloe vera* was demonstrated to control serum ALT level in HFFD treated mice. This is an indication that *Aloe vera* could prevent liver in injury that occur as a result of fat rich diet consumption. The possible mechanism by which *Aloe vera* prevent liver injury could be either through preventing hepatocytes damage and/or enhancing hepatocytes function.

Also the present study reports decrease in albumin and total protein concentration following HFFD consumption. Previous studies reported fatty acids to affect albumin’s redox status, enhance Cys34 oxidation and promoting lipid peroxidation leading to the development of metabolic disorders [25]. In the present study, *Aloe vera* improved the albumin and total protein concentration of HFFD fed mice. Albumin constitute about 20% of the proteins synthesized by hepatocytes in the liver and accounted for about 50% of the total plasma proteins [26]. Therefore, the increase in total protein concentration that was observed in the present study might be due to albumin upsurge. Serum albumin level is correlated with the number of normal/functional hepatocytes. Hence, hypoalbuminemia is associated with hepatocytes dysfunction and increase mortality rate [27,28]. *Aloe vera* might have prevent hepatocytes degeneration and enhance its function to produce more albumin. Hence, the normal histology that was observed in the liver of HFFD+AV treated mice in the present study.

Previous studies demonstrated that consumption of diet rich in fructose and fat is associated with oxidative stress and inflammation [29,30]. The decrease in catalase and superoxide dismutase activities that was observed in HFFD fed mice in the present study also demonstrated HFFD induced oxidative stress. *Aloe vera* supplementation enhanced the activities of catalase and superoxide dismutase of HFFD treated mice in the present study. Earlier report showed that Aloe vera increases antioxidant status and prevent oxidative stress in rodents [22,31,32]. The possible mechanism through which *Aloe vera* elicits antioxidant activity could be through scavenging free radicals or elevating serum albumin level as observed in the HFFD=AV treated mice in the current study. Albumin accounts for about 80% of extracellular thiols giving it the ability to scavenge free radical and prevent oxidative stress [33]. Albumin also regulate oncotic pressure and transport various ligands including ions, bilirubin, fatty acids and drugs. The role of albumin in oncotic pressure regulation is related to its high extracellular concentration and net negative charge [34]. Therefore, the antioxidant activity of *Aloe vera* is might be due its ability to enhance albumin production.

## Conclusions

*Aloe vera* was shown to protect against HFFD induced oxidative stress, hyperlipidemia and liver dysfunction. These findings could provide a clue for further research on food supplements for preventing oxidative stress related diseases and metabolic disorders in humans. However, high fat and fructose consumption should be reduced to the barest minimum since *Aloe vera* supplementation did not completely prevent lipid peroxidation.

## Acknowledgments

None

